# Frame Rate Up-Conversion in Echocardiography Images, Using Manifold-Learning and Image Registration

**DOI:** 10.1101/407072

**Authors:** A. Daneshi, H. Behnam, Z. Alizadeh Sani

## Abstract

In this paper, we propose a new temporal frame interpolation algorithm for frame rate up-conversion (FRUC) in echocardiography images. This algorithm employs a combination of dimension reduction techniques and image registration to increase frame rate.

If the distance between two successive frames of a video be great, motion jerkiness will appear between them and visual quality of the video will decrease. Some parts of heart have a very high speed motion, and echocardiography videos, obtained by available systems can’t take enough number of frames to show them well. So, to achieve an echocardiography image set with a better visual quality, more frames are necessary between two frames at a great distance. Here, we use dimension reduction techniques to find out the number of suitable frames between two consecutive frames to show the fast motions better, but don’t take much time. We project images to a 3-dimentional space by this way. Greater difference between the frames, results greater distance between corresponding embedded points. Thus, the distance between the embedded points is a scale for the suitable number of frames, needed between two successive frames. These frames are produced with the registration techniques.

On the other hand, heart doesn’t have a constant speed during a cycle, but echocardiography images are recorded with constant speed. So, frames at a greater distance show fast motions of the heart, and frames at a lower distance show slow motions of the heart. While, we put unequal number of frames between successive frames, and in this way remove temporal coordination of the image set. To solve this problem, we put efficient number of linear average of available frames, in places that the number of inserted frames in between available frames is less than maximum to obtain an equal number of frames between all successive frames.

## 1. Introduction

Echocardiography has become the predominant imaging modality in diagnostic cardiology, because it is noninvasive, inexpensive, and able to show moving parts in real-time. However, transient small motion of the heart wall and valves can’t be shown properly using low frame-rate imaging techniques. High frame rate is especially important in studying intra-cardiac structures where valve closure times are on the order of micro seconds. Current of-the-shelf ultrasound imaging systems have a limited frame rate because of the time it takes to send and receive all of the ultrasound beams necessary to reconstruct an image. While this frame rate is sufficient for real time human observation of basic ventricular function and assessment valve ability, understanding cardiac dynamics requires greater frame rates.

Several alternative methods have been developed to increase the ultrasound frame rate such as coded-excitation ultrasound imaging [1]–[7] and parallel processing techniques [8]–[10]. Some others increased the frame rate by reducing the size of the field of view and the total number of beams [11], [12]. The echocardiogram (ECG)-gating technique in ultrasound imaging is another method, recently introduced [13]. It divides the total field of view to seven equal sectors, and takes ECG signal and echocardiography images of each sector simultaneously. Then, combines the individual sectors into a large field-of-view at high beam density and also attains high frame rates. This last method assumes all heart cycles during a breath are the same. Also, because the respiratory motion could affect the heart’s position, breath-holding during the entire scan was required for higher composite imaging performance. Another is the digital scan converter (DSC) introduced by Chang et al. [14]. In this technique, a sparse beam array is send to the heart and a low quality image is produced in a short time. Then, a number of pixels are interpolated between available pixels. This method is useful for the small cases, such as mouse, which need a small depth of penetration.

Frame rate up-conversion (FRUC) is to increase the frame rate of a video by interpolating new frames and inserting them in between consecutive frames. Generally, FRUC can be classified into two groups. The first group interpolates the skipped frame along temporal axis without taking the object motion into account. Methods such as frame repetition (FR) and frame averaging (FA) belong to this group. These algorithms are very simple; but, produce “jerkiness” into the motion portrayal and blurriness on object boundaries [15]. The second group, named motion-compensated interpolation (MCI), interpolates the skipped frames along motion trajectory exploiting motion information between current successive frames [16]. Second group is more accurate than first group.

Fujiwara and Taguchi proposed a MCI method based on block matching algorithm (BMA). Since the property of the BMA is changed by the size of the blocks, it is desirable for small moving objects to set block size small, and in global motion region to set the block size large. Fujiwara and Taguchi used multi-size blocks and obtained less block artifact and more clear images [176].

Thaipanich and Wu proposed a MCI method for the cases that input video has abrupt illumination change and/or a very low frame rate [18]. However, because echocardiography images don’t have sudden changes in illumination and also satisfy a low threshold for frame rate, this method is not practical in these images.

We present a low complexity technique for exploiting the relationship between successive frames using the manifold learning algorithm and then interpolating new frames between available frames using MCI and FA.

The remains of this paper are organized as follows. The section 2 gives a mathematical background of the manifold learning algorithm and registration technique used here. Section 3 shows the results and section 4 presents the discussions. Finally, section 5 concludes this paper.

## 2. Materials and Methods

### 2.1 Materials

Used Manifold learning Algorithm (LLE)

Manifold learning algorithms attempt to expose intrinsic parameters in order to find a low-dimensional representation of the data.

Suppose that the original data consists of *n* data samples (observations) of the *X* data-set with dimension *D*. The Locally Linear Embedding (LLE) algorithm projects these observations into a new data-set *Y* consisting of *n* points with dimension *d* (where *d* <*D* and often *d* ≪*D*), while retaining the geometry of the data such as possible:

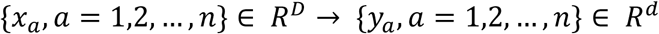

The embedding is optimized to preserve the local configurations of nearest neighbors. Keeping geometry of data is important in our proposed method, so, LLE algorithm is suitable here.

As shown in figure (1), LLE can be represented in three main steps [19]:

**Figure 1:**
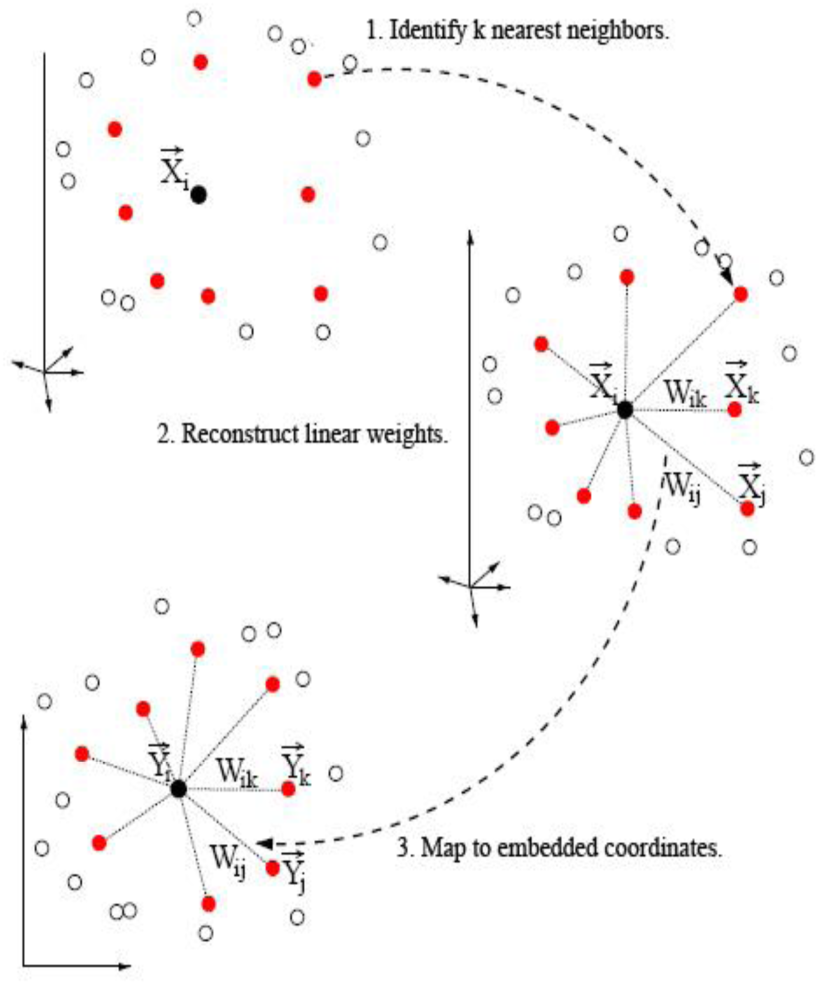
Summary of the LLE algorithm, mapping high dimensional inputs 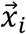 to low dimensional outputs 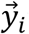 via local linear reconstruction weights W_*ij*_. (The image is reproduced from [19]).

Identify the *k* nearest neighbors for each data point. This can be done in two ways. (a) Having a constant *k*, find the *k* nearest neighbors as measured by Euclidian distance. (b) Determine data points which have a determined distance (radius of neighborhood) from that specific data point.

Model the manifold as a collection of linear patches and attempt to characterize the geometry of these linear patches. To do so, attempt to represent *x_i_* as a weighted, convex combination of its nearest neighbors. These linear weights *w_ij_*, must be chosen to minimize the following cost function

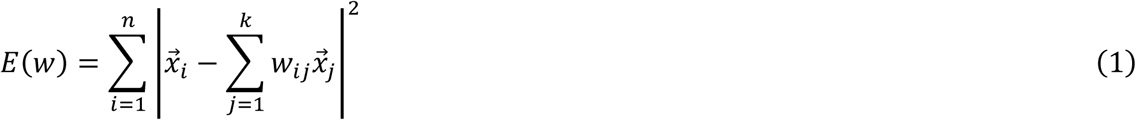

The weight matrix Wis used as a surrogate for the local geometry of the patches. Intuitively, *W_i_* reveals the layout of the points around *x_i_*. There are a couple of constraints on the weights: each row must sum to one 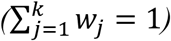 equivalently, each point is represented as a convex combination of its neighbors, and *w_ij_* = 0 for *j* < *k*. The second constraint reflects that LLE is a local method; the first makes the weights invariant to global translations: if each *x_i_* is shifted byα, then

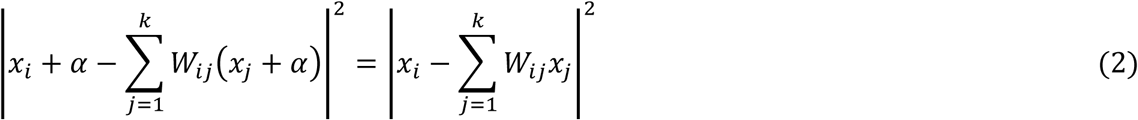

So *W* is unaffected. Moreover, *W* is invariant to global rotations and scalings. Fortunately, there is a closed form solution for *W*, which may be derived using Lagrange multipliers. In particular, the reconstruction weights for each point *x_i_* are given by

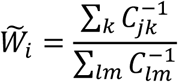

Where *C* is the local covariance matrix with entries 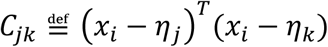, and *η_j_* (and) *η_k_* are neighbors of 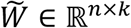 is then transformed into the sparse *n* × *n* matrix 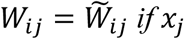 is the *l*th neighbor of *x_i_*, and is 0 if *j* ∉ *N*(*j*)

*W_i_* is a characterization of the local geometry around *x_i_* of the manifold. Since it is invariant to rotations, scaling, and translations, *W_i_* will also characterize the local geometry around *x_i_* for any linear mapping of the neighborhood patch surround *x_i_*. We expect that the chart from this neighborhood patch to the parameter space should be approximately linear since the manifold is smooth. Thus, *W_i_* should capture the local geometry of the parameter space as well. The next step of the algorithm finds a configuration in d-dimension (the dimensionality of the parameter space) whose local geometry is characterized well by *W*. *d* must be known a priori or estimated.

Learn embedding which preserves the reconstruction weights. In this step, each low dimensional output *y_i_* is mapped by a high dimensional input xi quoting global internal coordinates on the manifold. This, however, is possible by selecting the d-dimensional coordinates of each output *y_i_* to minimize the following cost function:

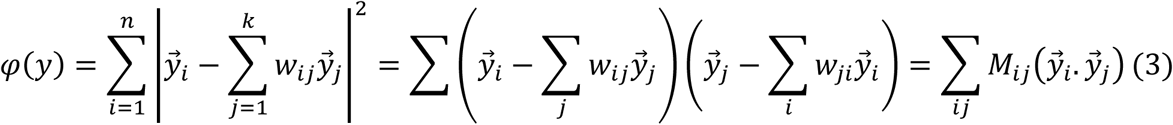

Where 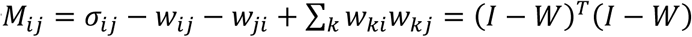

There are a couple of constraints on *Y*. First, *Y^T^Y*= I which forces the solution to be of rank *d*. Second, ∑_i_ *Y_i_*= 0; this constraint centers the embedding on the origin. This cost function is a form of Rayleigh’s quotient, so it is minimized by setting the columns of *Y* to the bottom d eigenvectors of *M*. However, the bottom (eigenvector, eigenvalue) pair is (1,0), so the final coordinate of each point is identical. To avoid this degenerate dimension, the d-dimensional embedding is given by the bottom non-constant d eigenvectors [19].

Here, in the first step, the *K* nearest neighbors for each data sample are determined as measured by the Euclidean distance. The LLE algorithm’s outcomes are normally stable over a range of neighborhood sizes [19]; in this study we chose *k* = 8 nearest neighbors for each data point.

While applying LLE algorithm on a specific image set, each pixel introduces one dimension. So, dimensionality of an image with *N* × *N* pixels is *N*^2^. That is a very large number. Making use of manifold learning, we can expose main parameters of echocardiography images and project them into a very low dimensional feature space.

Registration technique

The goal of image registration is to find the optimal transformation *T*, which will map any pixel in the floating image *I_F_*(*X,Y*) to its corresponding pixel in the reference image *I_R_*(*X,Y*) Floating image should take geometry of reference image, retaining intensity of pixels. To solve this problem, one can consider a number of points in each image, and find their correspondence to each other. Then the problem is to find a mathematical equation which projects these points in the floating image into their corresponding points in the reference image.

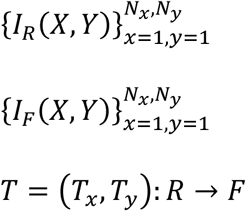

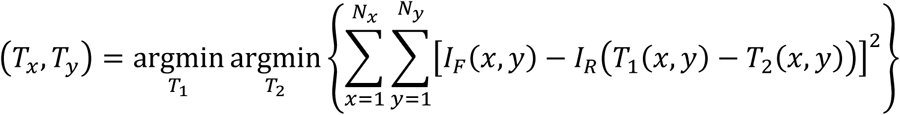

In above formulas, *I_F_*(*X,Y*) is the floating image *I_R_*(*X,Y*) is the reference image, *N_x_* and *N_y_* denote dimensions of the image, and *T*= (*T_x_*, *T_y_*) is the transformation that projects the floating image into the reference image. In reality, image registration transforms the floating image among a number of motion vectors (MVs) to produce an image with the most possible similarity to reference image.

The result of registration is an image which intensity values of its pixels are the same as the first image and its geometry is like the second image.

### 1. Experiment

The 2-D apical four chamber image sequences of eight healthy volunteers and two patients (endocardit and prosthetic mitral valve) are acquired using an ultrasound machine (General Electric, vivid 3) with a 1.7 MHz probe, including the ECG recording Echocardiography images are recorded at Rajaie Cardiovascular Medical and Research Center. We processed our method using MATLAB (MathWorksInc, USA) with a standard laptop computer (2 GHz Pentium, 4 GB RAM).

LLE algorithm is executed on these ten volunteer cases echocardiography images. Frame-rates and heart-rates of these cases are shown in table (1). Each frame had 401*461 pixels. Using LLE algorithm with k=8 nearest neighbors, a number of consecutive frames were projected into a three-dimensional space called “feature space”. Figures (2) show the image manifolds of one of the normal cases and endocardit case. Each “*” sign remarks one frame embedded in three-dimensional space.

**Figure 2:**
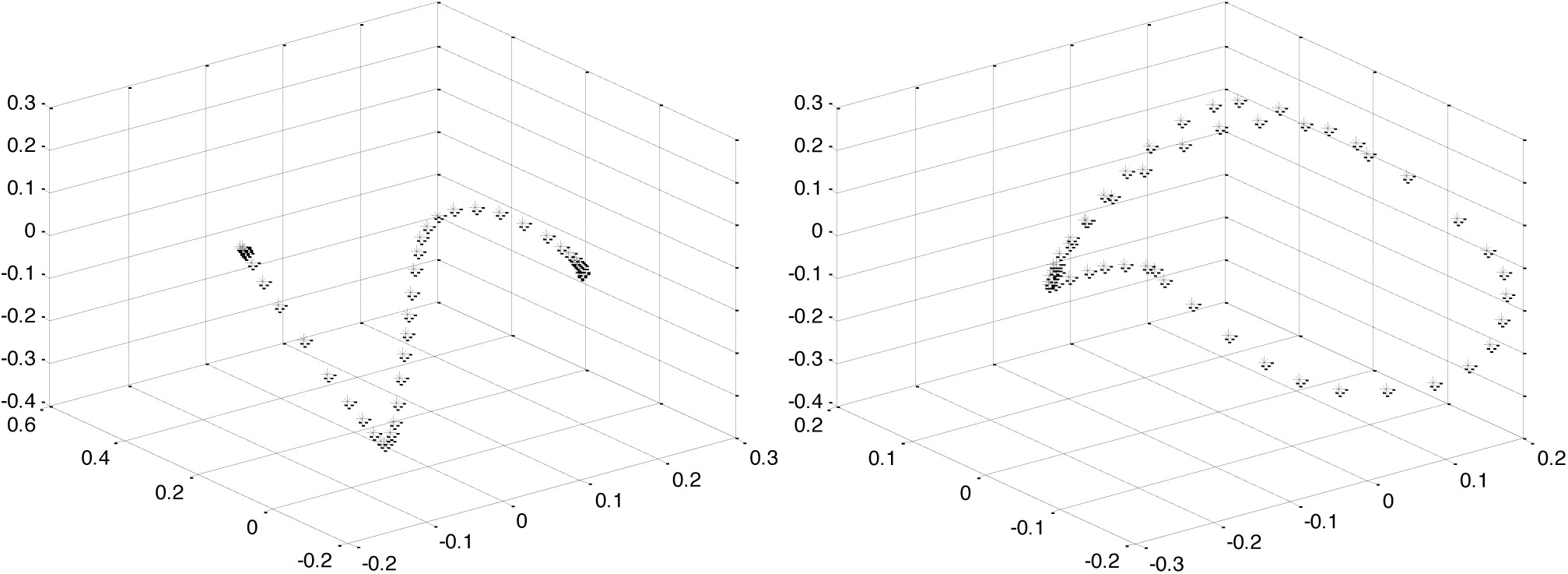
left: The three dimensional non-linear embedding of 50 frames of a normal heart (case 1) using LLE algorithm (k=8). Right: The three dimensional non-linear embedding of 50 frames of a heart with endocardit disease using LLE algorithm (k=8).

We set the dimension of the feature space equal three, because this is the least dimensionality that can catch all of the important elements making difference between frames.

The relationship of two consecutive images was retained during embedding; so, the distance between two successive “*”s was a criterion of the difference between corresponding frames. The average of these lengths was calculated and all of them were divided into this value and rounded to the nearest integer less than or equal to. The result was called characteristic number. The maximum characteristic number showed the suitable number of frames to be inserted among original frames.

Image registration was used where the division and rounding operations yielded a nonzero value. In this cases, motion vectors (MVs) were calculated and cut into a number of smaller vectors with equal length, named as chopped vectors. Figure (3) illustrates this for the case that difference of two successive images was twofold of the mean difference.

**Figure 3:**
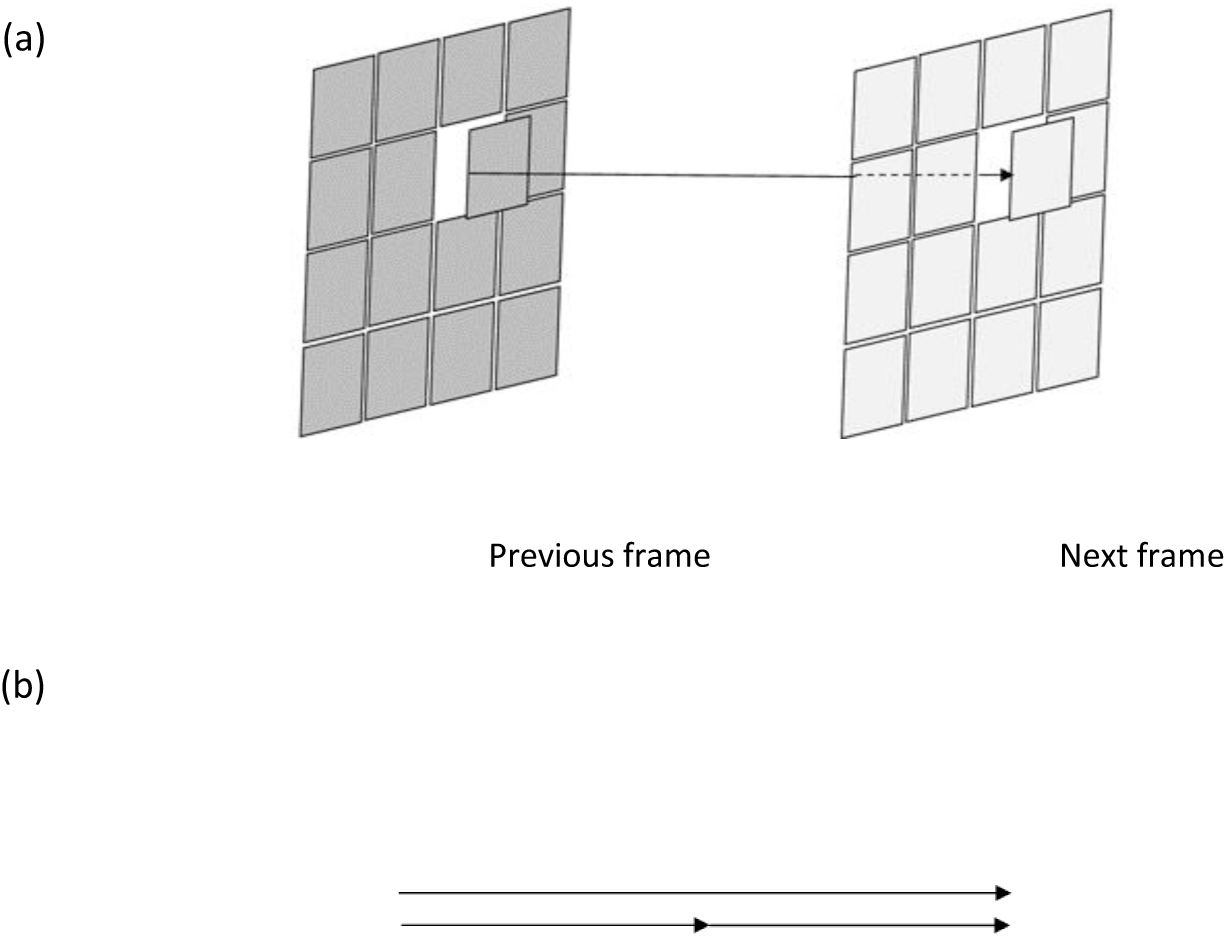
(a) Two successive frames and one of the motion vectors between them. (b) Sample motion vector divided to two vectors.

Considering two frames of the cycle, first frame was added with chopped vectors to construct middle frames. Since the number of chopped vectors between each pair of frames was different, to retain timing properties of the cycle, we used linear average of available frames to reach an equal number of frames between each pair of original successive frames. Using registration to produce new frames between two successive frames with a small difference takes a lot of time, but don’t give much more information than averaging. So, we don’t use registration for all frames.

First, we found out the difference between each characteristic number and the maximum characteristic number. Then, we used below algorithm.

Since maximum characteristic number of echocardiography images was less than three, there was four possible difference numbers. When the difference was zero, linear averaging wasn’t applied and all of unoriginal frames were produced using registration algorithm. When the difference was one, we inserted linear average of first frame and first registered frame between these two frames. When the difference was two, we had one registered frame, and inserted the linear average of this frame and two original frames before and after it. When the difference was three, we had no registered frame. So we inserted linear average of the two original frames between them; then, inserted average of this new frame and original frames before and after it.

Figure (4) illustrates this for the case that maximum characteristic number was three. Applying above algorithm on 50 frames of normal cases and endocardit and prosthetic mitral valve cases resulted different number of frames, shown in table (1).

**Figure 4:**
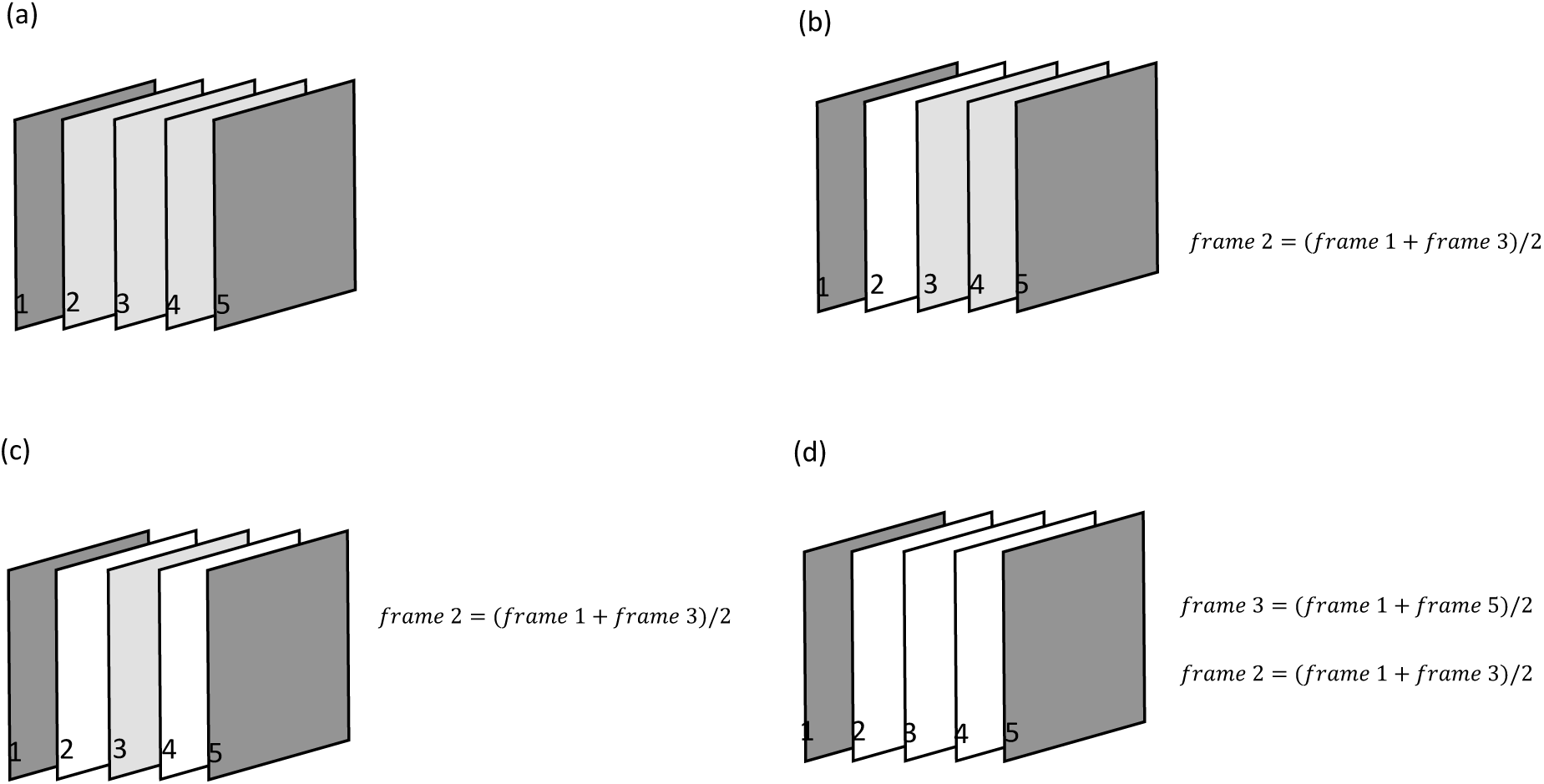
(a) difference= zero. (b) difference= one. (c) difference= two. (d) difference= three.

## 3. Results

Information about original echocardiography images and the frame rate up converted images is shown in table (1). To obtain proper frame rate, the number of result frames should be divided to the number of embedded frames and multiplied to the original frame rate. But, frame rates higher than 100 frames per second need special software and hardware.

Since maximum characteristic number of echocardiography cycle was less than three, and number of original frames used evaluate presented algorithm was 50, the number of result frames is less than 200.

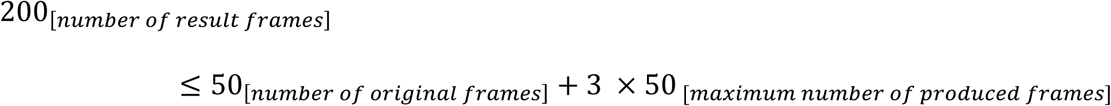

To evaluate registration algorithm, half of the frames of a normal cardiac cycle were removed. Indeed, even indexed frames were removed. Then, using mentioned registration algorithm, eliminated frames were reproduced. Figure (5) shows some original and reproduced frames and Figure (6) shows the squared difference values and the normalized cross correlation values between the first 50 even indexed original and reproduced frames.

**Figure 5:**
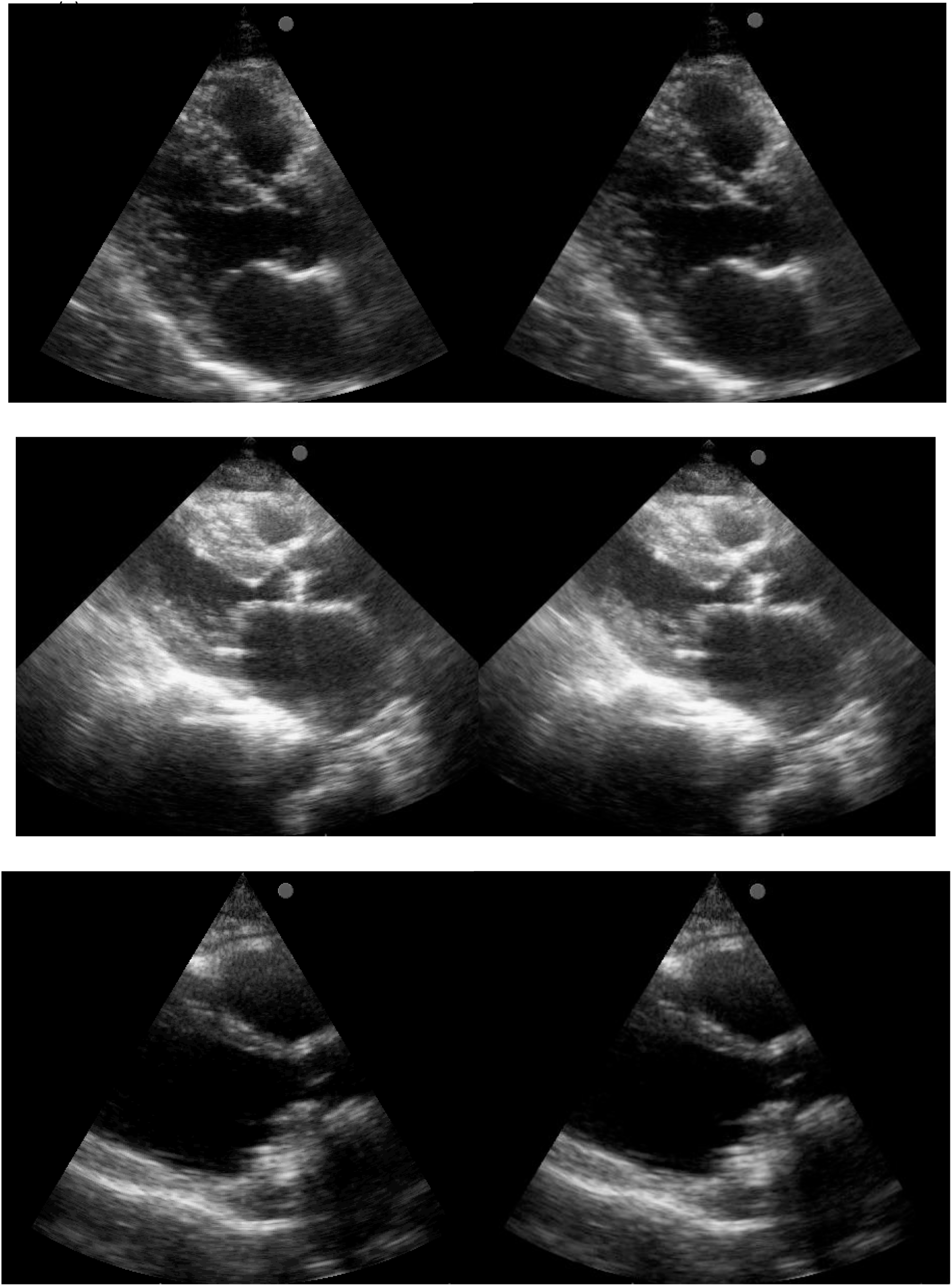
(a) One frame of the normal heart echocardiography. Left: original frame, Right: reproduced frame. (b) One frame of the endocardit case echocardiography. Left: original frame, Right: reproduced frame. (c) One frame of echocardiography of the heart with prosthetic valve. Left: original frame, Right: reproduced frame.

## 4. Discussion

Frames shown in figure (5) are original and reproduced frames using registration technique only. Produced videos, using proposed algorithm are available here.

Figure (6) shows the squared difference values and the normalized cross correlation values between the first 50 even indexed original and reproduced frames. It can be observed that squared differences are very low (close to zero) and the normalized cross correlation is very close to one. So, the frames obtained by mentioned registration algorithm, are very similar to original eliminated frames.

**Figure 6:**
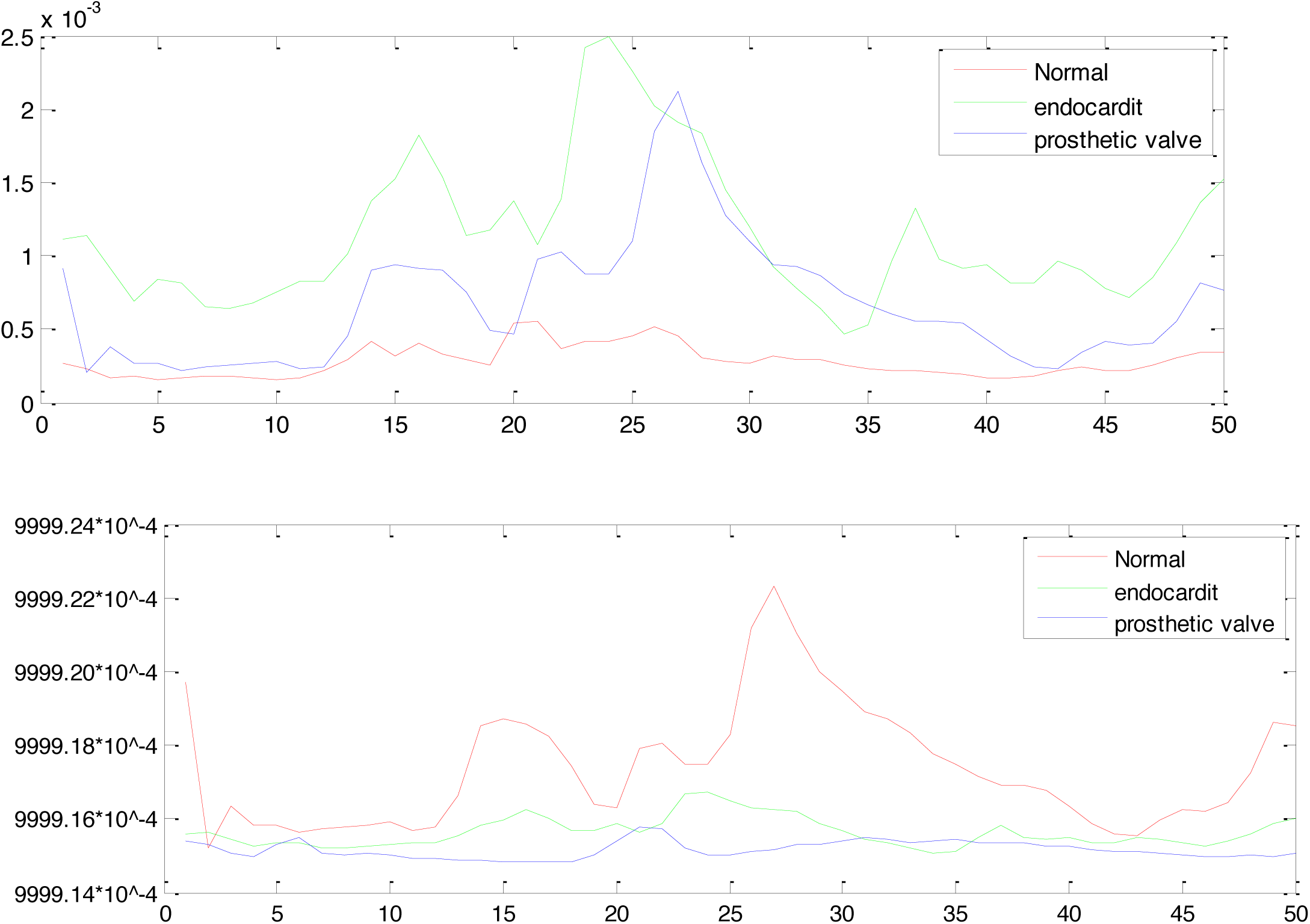
(a) the squared difference values between the first 50 even indexed original and reproduced frames, (b): the normalized cross correlation values between the first 50 even indexed original and reproduced frames.

As shown in table 1, frame rate of original echocardiography sets is usually more than 40 (f/s). So, increasing the number of frames, it is also ordinary to increase the show rate, the same number of times, to achieve an echocardiography set with a rate familiar for the heart specialists. For example, if the frame rate of the original set is 40 (f/s), and the number of frames in the new set is threefold of that in the original set, the frame rate of the new set should be 120 (f/s). However, with the available monitors it is not possible to show frames with a rate more than 100 (f/s). But nonetheless, heart specialists could see more details in echocardiography sets developed with the algorithm proposed in this paper. They do not need to resize echocardiography window to survey valves and other small parts of the heart. In figure 4, green circles show one of the most important parts for diagnosis. It is clear that produced frame shows this part much better.

**Table 1:** Total information about the three cases participated in this research.

|  | *Heart Rate (BPM)* | *Frame Rate (f/s)* | *Number of Result Frames* |
| --- | --- | --- | --- |
| *Case 1 (Normal)* | 62 | 82 | 127 |
| *Case 2 (Normal)* | 71 | 43 | 175 |
| *Case 3 (Normal)* | 71 | 64 | 126 |
| *Case 4 (Normal)* | 128 | 77 | 125 |
| *Case 5 (Normal)* | 74 | 68 | 177 |
| *Case 6 (Normal)* | 72 | 68 | 177 |
| *Case 7 (Normal)* | 68 | 53 | 177 |
| *Case 8 (Normal)* | 91 | 46 | 121 |
| *Case 9 (Endocardit)* | 76 | 46 | 120 |
| *Case 10 (Prosthetic valve)* | 75 | 79 | 178 |

## 5. Conclusion

In this paper, a low complexity FRUC method for echocardiography images is proposed. Our method finds the motion trajectory of echocardiography images and the relationship between consecutive frames. Then uses image registration techniques to find motion vectors among successive frames. Considering these motion vectors, a number of frames are inserted between available frames. At last, to retain timing properties of the image set, some original frames are repeated and the number of inserted frames between each pair if original frames is set equal.

